# Visual working memory representations in visual and parietal cortex do not remap after eye movements

**DOI:** 10.1101/747329

**Authors:** Tao He, Matthias Ekman, Annelinde R.E. Vandenbroucke, Floris P. de Lange

## Abstract

It has been suggested that our visual system does not only process stimuli that are directly available to our eyes, but also has a role in maintaining information in VWM over a period of seconds. It remains unclear however what happens to VWM representations in the visual system when we make saccades. Here, we tested the hypothesis that VWM representations are remapped within the visual system after making saccades. We directly compared the content of VWM for saccade and no-saccade conditions using MVPA of delay-related activity measured with fMRI. We found that when participants did not make a saccade, VWM representations were robustly present in contralateral early visual cortex. When making a saccade, VWM representations degraded in contralateral V1-V3 after the saccade shifted the location of the remembered grating to the opposite visual field. However, contrary to our hypothesis we found no evidence for the representations of the remembered grating at the saccadic target location in the opposite visual field, suggesting that there is no evidence for remapping of VWM in early visual cortex. Interestingly, IPS showed persistent VWM representations in both the saccade and no-saccade condition. Together, our results indicate that VWM representations in early visual cortex are not remapped across eye movements, potentially limiting the role of early visual cortex in VWM storage.

**Highlights:**

- Visual working memory (VWM) representations do not remap after making saccades
- Eye movement degrade VWM representations in early visual cortex, limiting the role of early visual cortex in VWM storage
- Parietal cortex shows persistent VWM representations across saccades

## Introduction

Research in the past decade has suggested that our visual system may not only process incoming information, but also be relevant for maintaining internal representations of previously observed visual stimuli, i.e. visual working memory (VWM). The ability to maintain information that is no longer in view is critical for reasoning about and mentally manipulating visual information despite temporal discontinuous in visual inputs that occur for example during eye movements, occlusions, and object motion (Curtis and D’Esposito 2003; Serences 2016; Lorenc et al. 2018).

VWM information has been observed both in early sensory areas (Ester et al. 2009; Harrison and Tong 2009; Albers et al. 2013; Sreenivasan et al. 2014), as well as higher-order parietal (Christophel et al. 2015; Ester et al. 2015; Bettencourt and Xu 2016), and prefrontal regions (Goldman-Rakic 1995; Durstewitz et al. 2000; reviewed in Riley and Constantinidis 2016). Moreover, VWM representations in early visual cortex have also been found to be spatially specific, maintained in a retinotopic manner, at least in V1 and V2 (Pratte and Tong 2014; although see Ester et al. 2009). Crucially however, in these previous studies, participants were instructed to maintain fixation while remembering a visual stimulus during a retention period. While this is a common approach to investigate VWM in a lab environment, it is arguably quite different from real-world settings that are marked by multiple eye movements per second. Humans perform about three saccadic eye movements per second in order to guide the fovea toward regions of interest in the visual field (Rayner 1998). Previous research has indicated that both stimulus features and attentional pointers are remapped across eye movement to aid visual stability (Melcher 2007; Rolfs et al. 2011; He et al. 2018). It is currently unclear however, whether VWM is also remapped within the visual system after making saccades.

One possibility is that VWM representations are dynamically remapped and follow the shift of the retinotopic location caused by eye movements. In this case, the cortical location representing the VWM would be updated after an eye-movement. Alternatively, if VWM does not remap, working memory information might be maintained across multiple brain regions following eye movements. On this account, after every eye movement, the up-to-date feature of the stimulus (e.g., the latest location of the stimuli) would be integrated with the previous working memory information.

Here, we tested the potential remapping of VWM in the early visual system, by presenting participants with an orientated grating in either the left or right visual field. The grating orientation had to be maintained for a subsequent retention period, while we measured blood oxygenation level-dependent (BOLD) signals with functional magnetic resonance imaging (fMRI). Using multivariate pattern analysis (MVPA), we attempted to decode the remembered orientation in both the contralateral and the ipsilateral hemispheres during trials in which participants performed either a saccade or maintained fixation during the working memory period.

To preview, we found a contralateral VWM representation in the early visual system. This VWM representation degraded when participants made a saccade and was not remapped to ipsilateral visual cortex. In contrast, VWM representations in IPS persisted after saccades. These findings suggest a limited role of the early visual system in VWM.

## Materials and Methods

### Data and code availability

All data and code used for stimulus presentation and analysis is freely available on the Donders Repository (https://data.donders.ru.nl/login/reviewer-77603154/28KfrmSvRmvGnYdOkGqdQQfMufTI7W29jY686hMMaro).

### Participants

Thirty-four healthy participants (18 females, mean age 24.3 years, ranging from 19 to 33 years) with normal or corrected-to-normal vision were recruited from the institute’s subject pool in exchange for either monetary compensation or study credits. This sample size of N = 34 included subjects ensured 80% power to detect an effect size of at least Cohen’s d ≥0.5. All participants were naive with respect to the purposes of the study. The experiments were approved by the Radboud University Institutional Review Board and were carried out in accordance with the guidelines expressed in the Declaration of Helsinki. Written informed consent was obtained from all participants prior to the start of the study. Only participants who completed the entire experimental protocol (i.e., the behavioral experiment and the two fMRI sessions) were included in the final analysis. Data from six participants were excluded: one participant didn’t complete the fMRI sessions due to discomfort, one participant failed to follow the instructions (i.e., fixate and saccade) during experiment, four participants had excessive head motion (motion cutoff = 1 mm).

### Stimuli

Stimuli were programmed in MATLAB (v2016a, The MathWorks, Inc., Natick, MA) using the Psychtoolbox (Brainard 1997). The circular sinusoidal grating stimulus subtended 10° and was centered with a small jitter (0.3°) on the screen center. The grating was full contrast, with a spatial frequency of 1 cycles/degree, a random phase and an orientation of either 25° or 115° (with a small jitter of 3°) from the horizontal axis. The contrasts of the edges of the grating were linearly attenuated over the distance from 4.5° to 5.0° radius. Two filled dots (0.5°, one green, one black) were presented at the periphery of the screen (6° left or right away from the center of the screen), which were served as the fixation dot and saccade target in the task. In the behavioral training session, stimuli were presented on a 24-inch flat-panel display (BenQ XL2420T, 1,920 × 1,080 resolution, 60 Hz refresh rate). In the fMRI sessions, stimuli were displayed on a rear-projection screen using an EIKI LC-XL100L (EIKI, Rancho Santa Margarita, CA) multimedia projector (1,024 × 768 resolution, 60 Hz refresh rate).

### Experimental design

#### Behavioral training procedure

Prior to the fMRI scan sessions, all participants completed a one-hour behavioral training session to familiarize themselves with the fMRI main task and to establish their individual orientation discrimination threshold, which served as an initial orientation difference of the gratings in the following fMRI sessions. The experimental design of the behavioral training task was exactly the same as during the fMRI main task, except that the delay period and the inter-trial interval (ITI) were shortened to 3 s to reduce the experimental time. Participants completed two to three blocks of 56 trials until a stable orientation discrimination threshold was obtained, during which the eye movements were continuously monitored by an Eyelink 1000 plus eye tracker. In addition, participants were familiarized with the localizer tasks and the retinotopic mapping procedures that were used in the fMRI session at the end of the behavior session.

#### fMRI main task

The experimental design (Figure 1a) was adapted from a well-known delayed orientation discrimination task used before (Harrison and Tong 2009). Each trial began with two filled dots (one green, one black) that were presented at the periphery of the screen (2 s). Participants were asked to fixate at the green dot through the trial. Two oriented gratings were flashed sequentially at the center of the screen (which could be in the left or right visual field of the participants, depending on the location of the green fixation dot) for 200 ms, respectively, with an inter-stimulus interval of 400 ms, and followed by a retro-cue (400 ms, “1” or “2”) to indicate which orientation of the grating should be remembered during the following delay period. The sequence of the orientations (25° and 115°) were randomly chosen on each trial. On 10 out of 28 trials (No-saccade condition) in each run, the two dots did not swap their locations, which indicated to participants that they needed to keep their fixation at the initial fixation location during the 10 s delay period. On the remaining 18 trials (Saccade condition) in the run, the green dot was shifted to the opposite side of the screen at the beginning of the delay period (2 s after the offset of the retro-cue). This instructed participants to move their eyes to the opposite location of the screen and maintain their fixation at the new position for the remaining 8 s. A probe grating was presented at the screen center after the delay period and participants were required to make a judgement of whether the probe orientation was rotated clockwise or count-clockwise relative to the grating they memorized during the delay period. Finally, feedback was provided and a black fixation dot instructed participants to move their eyes to the center of the screen in preparation for the next trial. Trials were separated by an ITI of 9.6 s. The central fixation dot changed its color from black to gray at the last second of the ITI to indicate that the next trial was going to start.

**Figure 1.**
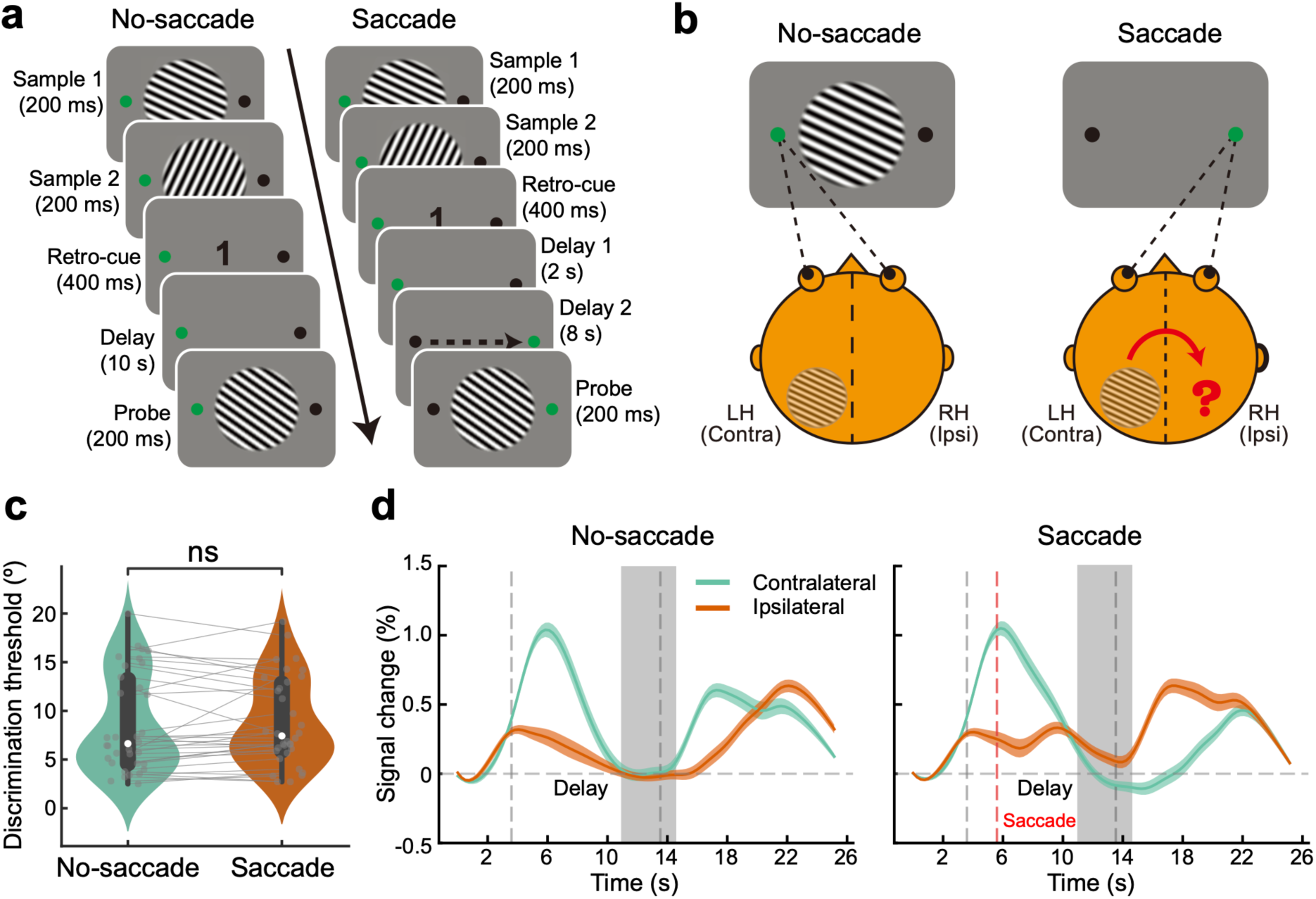
Experimental paradigm, behavioral performance and BOLD activity in the early visual cortex. **(a)** Participants performed a delayed orientation discrimination task. At the start of each trial, two dots were presented at the periphery of the screen and participants were instructed to fixate at the green dot. Two orientated gratings (25° and 115°) were successively flashed in the center of the screen, followed by a retro-cue (1 or 2) that indicated which grating to remember in the following delay period. Crucially, during a 10 s delay period, if the green dot shifted to the opposite side of the screen (*saccade condition*), participants had to make an eye movement to the opposite side and maintained at the new location through the following time period (the dashed arrow is for illustration only, not present in the actual test). Conversely, in the no-saccade condition, the green dot did not change its position and participants maintained fixation at the initial side throughout the entire trial. After the delay period a probe was presented, and participants indicated whether the probe was tilted clockwise or counterclockwise relative to the remembered grating. **(b)** Illustration of the experimental design probing remapping. In the no-saccade condition (left), the remembered grating presented in right visual field is initially represented in the left (contralateral) early visual cortex. In the saccade condition (right), after participants saccade from the left to the right side of the screen, we set out to examine whether VWM contents that initially generated in contralateral (left) early visual cortex transfer to the ipsilateral (right) visual cortex. **(c)** there was no difference in the mean discrimination threshold between trials with and without a saccade, demonstrating that the VWM performance was not impaired by the eye movement during the delay period. Grey dots with connecting lines denote individual participants. Colors are estimated densities, white dots are group medians, boxes are quartiles and whiskers are 1.5 interquartile range. **(d)** Group averaged (N = 34) BOLD time course in contralateral and ipsilateral early visual areas (V1 – V3). In the no-saccade condition, the presentation of grating stimuli evoked a higher BOLD response in contralateral than in ipsilateral early visual areas. In the saccade condition, while the BOLD activity was still higher in contralateral relative to ipsilateral early visual areas after presentation of the grating stimuli, the pattern inverted after presentation of the probe – BOLD activity became higher in ipsilateral than in contralateral early visual areas. The first vertical dashed gray line represents the onset of the gratings, while the second vertical dashed gray line indicates the onset of the probe. The vertical dashed red line in saccade condition indicates the onset of the saccade. The vertical gray bar represents the delay period (7.2 s - 10.8 s after onset of maintenance) that selected for multivariate analysis. BOLD activity data was interpolated and smoothed for display only, all statistical tests were applied before data interpolation. Shaded areas denote SEM.

In order to incentivize participants to form spatially specific visual working memories, four catch trials were included in each run. Stimuli and timing of the catch trials were identical to the main trials, except for the following changes. During the presentation of the probe in the catch trial, the probe was horizontally shifted 1.2° to the left or right with respect to the location of the sample gratings. Participants were instructed to not respond when the location of the probe grating did not match with the sample gratings. The rationale of this was that it forced participants to memorize the orientation stimulus at its original location in order to successfully perform the task, resulting in a spatially specific VWM representation.

Each run comprised 28 trials, consisting of 10 no-saccade trials (including 2 catch trials), 18 saccade trials (including 2 catch trials). We chose these trial numbers because the saccade trials were of primary interest and we therefore wanted to have maximal sensitivity for this condition. The order of trials was a pseudo-randomized within each run. Each trial lasted 26.4 s, consisting of a 16.8 s task period and a 9.6 s ITI. Each run lasted 12.32 min and started with 6 s of fixation that was discarded from the analysis.

### Staircase Procedure

The staircase procedure was used to ensure equal task difficulty across participants and to equate task difficulty levels across no-saccade and saccade conditions within participants. In the behavioral training session, immediately after the delay period, participants were asked to compare the tilted orientation between the probe and the internal remembered grating. The staircase procedure estimated the difference between probe and remembered grating that ensured 75% performance, using QUEST (Watson and Pelli 1983; Prins and Kingdom 2018). A maximum orientation difference of 20° between the probe and remembered grating was enforced. The staircase was initiated with an orientation difference of 10° and dynamically adapted according to participants’ performance on previous trial until a stable threshold was acquired. This threshold was used as a seed in the following fMRI sessions and the same staircase procedure was also used during scanning.

### Eye tracking

Eye position was monitored with an MR-compatible Eyelink 1000 (SR Research Ltd., Ottawa, Canada) eye tracker. Only the left eye was recorded in the scanner. Pupil and corneal reflection were sampled at 1000 Hz and analyzed offline to ensure that participants fixated at the correct location. The eye tracker was calibrated at the beginning of each session and repeated between runs if necessary. In 10 out of 192 runs, the eye tracker signal was lost during scanning due to subjects’ head motion or technical problems. During these runs, the experimenter monitored the eye position online via the live video feed from the camera. All participants were trained on the fixation task and to perform saccades in the behavioral training session prior to the scanning session.

### fMRI parameters

Functional and anatomical images were carried out with a 3T Siemens Prisma fit MRI system (Siemens, Erlangen, Germany), using a 32-channel headcoil. Functional images were acquired using a whole brain T2*-weighted multiband-4 sequence (TR/TE = 1,200/39 ms, voxel size 2.4×2.4×2.4 mm, 56 transversal slices, 65° flip angle, A/P phase encoding direction, FOV = 210 mm, BW = 2030 Hz/Px). Anatomical images were acquired using a T1-weighted magnetization prepared rapid gradient echo (MP-RAGE) sequence (TR/TE = 2,300/3.03ms, voxel size 1×1×1 mm, 192 transversal slices, 8° flip angle).

### fMRI Preprocessing

fMRI data were preprocessed using FSL (FMRIB Software Library; Oxford, UK; www.fmrib.ox.ac.uk/fsl; Smith et al. 2004, RRID:SCR_002823), including motion correction (six-parameter affine transform), temporal high-pass filtering (100 s) and Savitzky–Golay low-pass filter (time window length = 11 TRs, polynomial order = 3; Savitzky & Golay, 1964) for each run separately. No spatial smoothing and slice timing correction was performed. All univariate and multivariate analyses were performed in native subject space with custom python code using Nibabel [https://doi.org/10.5281/zenodo.1464282], Scipy (Oliphant 2007; Millman and Aivazis 2011) and scikit-learn (Pedregosa et al. 2011).

### Functional localizers

In addition to the main experiment, participants underwent two localizer runs, which were used to select voxels that maximally responded to stimuli presented in the contralateral hemifield in both the univariate and multivariate analysis. The same grating stimulus parameter were used in localizer runs as those in the main experiment. Participants fixated at the left or right green dot and the grating was presented at the center of screen for 16 s, with a frequency of 4 Hz. Throughout the localizer, participant had to fixate at the green dot and monitor a sequence of rapidly changing letters just above fixation, to which they had to respond by button press whenever a target letter (“X” or “Z”) occurred in a stream of non-target letters (“A”, “H”, “R”, “N”, “T”, “V”, “U”, “Y”). Letters were presented at a frequency of 2 Hz. When the green dot shifted from the left to the right side of the screen (or vice versa) at the end of the trial, participants had to move their eyes to follow the fixation dot. Each trial was separated by a 2 s ITI in which only the fixation cross was presented, to give participants enough time to saccade and stabilize their eyes in the new position. Two blocks of 20 trials (10 trials per visual field: left or right) were collected.

Images of the localizer runs were also preprocessed following the same procedure as the main tasks. Onsets and duration of the localizer blocks were convolved with a double gamma haemodynamic response function (HRF) and fitted using a general linear model in left and right hemisphere separately. For each participant a two-sided t-contrast was calculated contrasting between left and right hemispheres. Resulting statistical maps were thresholded at Z > 3.1 and a corrected cluster significance threshold of *p* < 0.05.

### Retinotopic mapping of early visual areas (V1 – V3)

Early visual areas (V1, V2, and V3) for each participant were identified using two retinotopic mapping blocks based on the standard traveling-wave method using rotating wedges (DeYoe et al. 1996; Engel et al. 1997; Wandell et al. 2007), consisting of a clockwise and a counterclockwise-rotating run. Participants were instructed to fixate at the center of the screen and fulfill rapid letter detection task (exact same as that in localizer runs). The BOLD responses to the wedges were used to estimate the polar angle of the visual field representation.

### Regions of interest (ROIs)

Early visual areas (V1 - V3) and the intraparietal sulcus (IPS0 – IPS5) within each hemisphere were independently chosen as the ROIs in our analyses. For the ROI of the early visual areas, we first used Freesurfer (http://surfer.nmr.mgh.harvard.edu/, Fischl et al. 2002) to define the gray–white matter boundary and perform cortical surface reconstruction. The borders of the early visual areas V1 - V3 were delineated based on retinotopic maps. The ROI of the intraparietal sulcus was defined based on anatomical probability maps of retinotopic areas in the intraparietal sulcus (IPS0 – IPS5) from the Probabilistic atlases (Wang et al. 2015). Finally, all surface-based ROIs were backward-transformed into participant’s native space. Next, the visually active voxels corresponding to the grating that was positioned at the left and right visual field was identified based on statistical activation maps from the functional localizer runs. Two types of ROI were defined in the decoding analyses: contralateral ROIs and ipsilateral ROIs. For instance, when decoding the orientation of the grating that was initially presented at participant’s right visual field, regardless of the no-saccade and saccade condition, the left hemisphere would be labeled as the contralateral ROIs, while the right hemisphere would be labeled as the ipsilateral ROIs (see Figure 1b) and vice versa. Finally, we selected the 150 most active voxels across visual areas V1, V2, V3, as well as 250 most active voxels across the combined early visual cortex (V1 - V3) and the entire intraparietal sulcus (IPS0 – IPS5) to perform the decoding analyses, separately.

### Univariate fMRI analyses

To estimate the BOLD response of each hemisphere in each condition, we selected the same voxels as those used for decoding. We separately modeled the onset of each trial for each hemisphere and for no-saccade and saccade condition within each run to fit voxel-wise general linear models (GLM) using FSL FEAT. An additional nuisance regressor of 24 motion regressors (FSL’s standard + extended motion parameters) were also added to the GLM. To quantify BOLD activity during the trial, contrasts between left and right hemisphere regressors for each condition were created. Multiple-comparison correction was performed using Gaussian random-field based cluster thresholding. The significance level was set at the cluster-forming threshold of z > 3.1 (i.e., *p* < 0.001, two-sided) and a cluster significance threshold of *p* < 0.05. All fMRI data were transformed from MRI signal intensity to units of percent signal change, calculated relative to the average level of activity for each voxel across the first volume of each trial within each run. BOLD activity over time were statistically tested using nonparametric cluster-based permutation t test (10,000 permutations; two-sided *p* < 0.05; cluster formation threshold *p* < 0.05) (Maris and Oostenveld 2007).

### Multivariate fMRI analyses

We used multivariate pattern analyses (MVPA) to determine whether the pattern of activity in each ROI and each hemisphere contained orientation information, as implemented in Scikit-learn 0.20.3 (Pedregosa et al. 2011). To this end, linear support vector machines (SVMs) were trained to discriminate between the two grating orientations based on the pattern of BOLD activity over voxels. In this study we used classification distance as an indication of the amount of orientation information being maintained in each hemisphere. To calculate the classification distance, we measured the distance of each sample to the separating hyperplane that was trained by a linear SVM with a positive/negative sign (the sign indicates the class). We then averaged the distance within each class and calculated the absolute distance between these two classes. While this approach can give us a binary predicted label (25° or 115°), which can be used to calculate classification accuracy, it additionally allows one to review the confidence of this classification and provide a continuous metric of the decoding performance. In each ROI, the larger the classification distance, the more confident the classifier is in determining the stimulus class based on the BOLD activity pattern, and hence the more orientation information is contained within the pattern of BOLD activity.

When training and testing within the main experiment we averaged the BOLD activity over time points 7.2 - 10.8 s after the onset of the delay period. This time period was selected to be as far from the onset of the two gratings at the beginning of the trial as possible, and therefore not reflect activity elicited by the stimuli, but also prior to the onset of the probe stimulus. These time series were then normalized on a voxel-by-voxel and run-by-run basis for each voxel using a z-scored transformation and sorted into one of sixteen bins based on four factors: hemisphere (left or right hemisphere), orientation (25° or 115°), stimulus location (left or right hemifield) and saccade condition (saccade or no-saccade). A leave-one-run-out cross-validation procedure was used to train the classifier where we trained in 7 (“training” dataset) out of 8 runs and tested on the remaining run (“test” dataset) for each hemisphere, stimulus location and saccade condition pair, separately. Within the independent “training” and “test” dataset, activation patterns comprising the mean response of each voxel during 25° or 115° trials were calculated. Finally, the decoding distances in each corresponding stimulus location and hemisphere pair were collapsed into the contralateral (i.e., left hemifield and right hemisphere, right hemifield and left hemisphere) or ipsilateral (i.e., left hemifield and left hemisphere, right hemifield and right hemisphere) hemisphere condition.

### Bayesian analyses

In order to further evaluate all the statistical tests, we performed the Bayesian equivalents of the above outlined analyses. JASP 0.10.2 (JASP Team, 2019, RRID:SCR_015823) was used to perform all Bayesian analyses, using default settings. Thus, for Bayesian t-tests a Cauchy prior width of 0.707 was chosen. Qualitative interpretations of Bayes Factors are based on criteria by Lee & Wagenmakers (2014).

## Results

Our primary goal was to investigate whether VWM representations in early visual cortex (V1-V3) are remapped after eye movements. We used MVPA (Kamitani and Tong 2005; Haynes and Rees 2006; Norman et al. 2006) to determine whether the information in VWM that was initially represented in the contralateral hemisphere (corresponding to the stimulus location) was remapped to the opposite (ipsilateral) hemisphere following an eye movement to the opposite visual field during the retention period. 34 participants performed a delayed orientation discrimination task, in which they initially fixated at the peripheral fixation dot while passively viewing two successive grating stimulus orientations (25° and 115°) in one visual field, followed by a cue and a delay period. Importantly, in 18 out of 28 trials (*saccade condition*) in each run, participants were asked to saccade to the opposite side of the screen during the delay period, while in the remaining 10 trials (*no-saccade condition*) in the same run, participants maintained fixation at the initial side through the entire trial. A probe grating was presented at the end of the trial (**Figure 1a, b**).

In order to encourage participants to memorize the grating in its original location, we also included catch trials in which the probe grating was horizontally shifted 1.2° with respect to the location of the sample gratings. Participants were instructed to discriminate the orientation between the probe and the memorized grating only when they were presented at the same location (see Methods for details). This experimental design was chosen to encourage participants to maintain both the orientation and location of the presented grating. Indeed, behavioral data showed that participants successfully withheld their response (i.e. correct rejection rate) at a rate of 63.6% (SD 18.37) and 62.87% (SD 19.58) in both no-saccade and saccade conditions when the probe was presented at a displaced location, and no difference was found between the two conditions (t_(33)_ = 0.257, *p* = 0.7989, Cohen’s *d* = 0.039; BF_10_ = 0.189), suggesting that participants remembered the orientation in a spatially specific way.

Behavioural data further confirmed that participants could successfully discriminate small differences between the cued grating and the probe grating (**Figure 1c**, mean discrimination threshold, no-saccade condition: 8.63°, SD 5.07; saccade condition: 8.89°, SD 4.55) at an accuracy of 82.1% (SD 5.05) and 81.1% (SD 4.43) in both no-saccade and saccade trials. Notably, there was no difference in accuracy between trials with and without a saccade (t_(33)_ = 1.125, *p* = 0.2687, Cohen’s *d* = 0.193; BF_10_ = 0.328), and no difference in the mean discrimination threshold (t_(33)_ = 0.66, *p* = 0.514, Cohen’s *d* = 0.113; BF_10_ = 0.225), indicating that the VWM performance was not impaired by the eye movement during the delay period.

A univariate BOLD analysis showed that the presentation of the grating stimuli induced stronger BOLD activity in contralateral than ipsilateral early visual areas (cluster permutation test, *p* = 0.0001). As expected, probe presentation at the end of the trial led to a BOLD activity increase in the contralateral hemisphere during no-saccade trials and a BOLD activity increase in the ipsilateral hemisphere, corresponding to the updated stimulus location after the eye movement, in the saccade condition (**Figure 1d**). Taken together, these results suggest that participants engaged successfully in the task.

In order to evaluate the potential remapping of VWM in early visual cortex, we assessed whether the patterns of activation in both hemispheres, contralateral and ipsilateral to the item location, contained the remembered orientation information during the delay period (**Figure 1d**, gray region). A leave-one-run-out cross-validation approach was used to train the classifier to discriminate the grating orientation (25° vs. 115°) within the working memory period and test on the left-out run. We used classification distance to measure the amount of orientation information present in the activity patterns (Dijkstra et al. 2019; Linde-Domingo et al. 2019). This approach results in a binary prediction (25° or 115°), but additionally yields a continuous metric of the decoding performance that can be seen as the confidence of the classification (see Methods for additional details).

### No remapping of VWM in early visual cortex following eye movements

During the no-saccade condition, activity patterns in the early visual areas (V1 - V3) contralateral to the grating location contained information about the maintained orientation during the delay period (7.2 s - 10.8 s after onset of maintenance) (**Figure 2**, dark green bar; t_(33)_ = 3.652, *p* = 0.0009, Cohen’s *d* = 0.63; BF_10_ = 35.369). There was also weak evidence for some orientation information in the ipsilateral hemisphere of V1 - V3 (**Figure 2**, dark orange bar; t_(33)_ = 2.192, *p* = 0.0356, Cohen’s *d* = 0.38; BF_10_ = 1.501), and no compelling evidence for a difference between the contralateral and ipsilateral hemisphere of V1 - V3 (t_(33)_ = 1.669, *p* = 0.1047, Cohen’s *d* = 0.32; BF_10_ = 0.642). The same pattern was also visible when inspecting the early visual areas separately (**Supplementary Figure 1**). However, in the saccade trials, in which participants made an eye movement that shifted the location of the remembered grating to the opposite visual field during the delay period, the classifier was not able to distinguish between grating orientations in V1 – V3 contralateral to the item location (**Figure 2**, light green bar; t_(33)_ = 0.03, *p* = 0.9765, Cohen’s *d* = 0.005; BF_10_ = 0.184), and similarly there was no compelling evidence for orientation information in ipsilateral V1-V3 (**Figure 2**, light orange bar; t_(33)_ = 1.905, *p* = 0.0655, Cohen’s *d* = 0.327; BF_10_ = 0.92). Again, there was no difference in orientation information between the contralateral and ipsilateral hemisphere of V1 – V3 (t_(33)_ = 1.722, *p* = 0.09438, Cohen’s *d* = 0.286; BF_10_ = 0.694). Crucially, there was a significant degradation of orientation information in the contralateral hemisphere of the early visual cortex after an eye movement (t_(33)_ = 2.748, *p* = 0.0096, Cohen’s *d* = 0.478; BF_10_ = 4.452).

**Figure 2.**
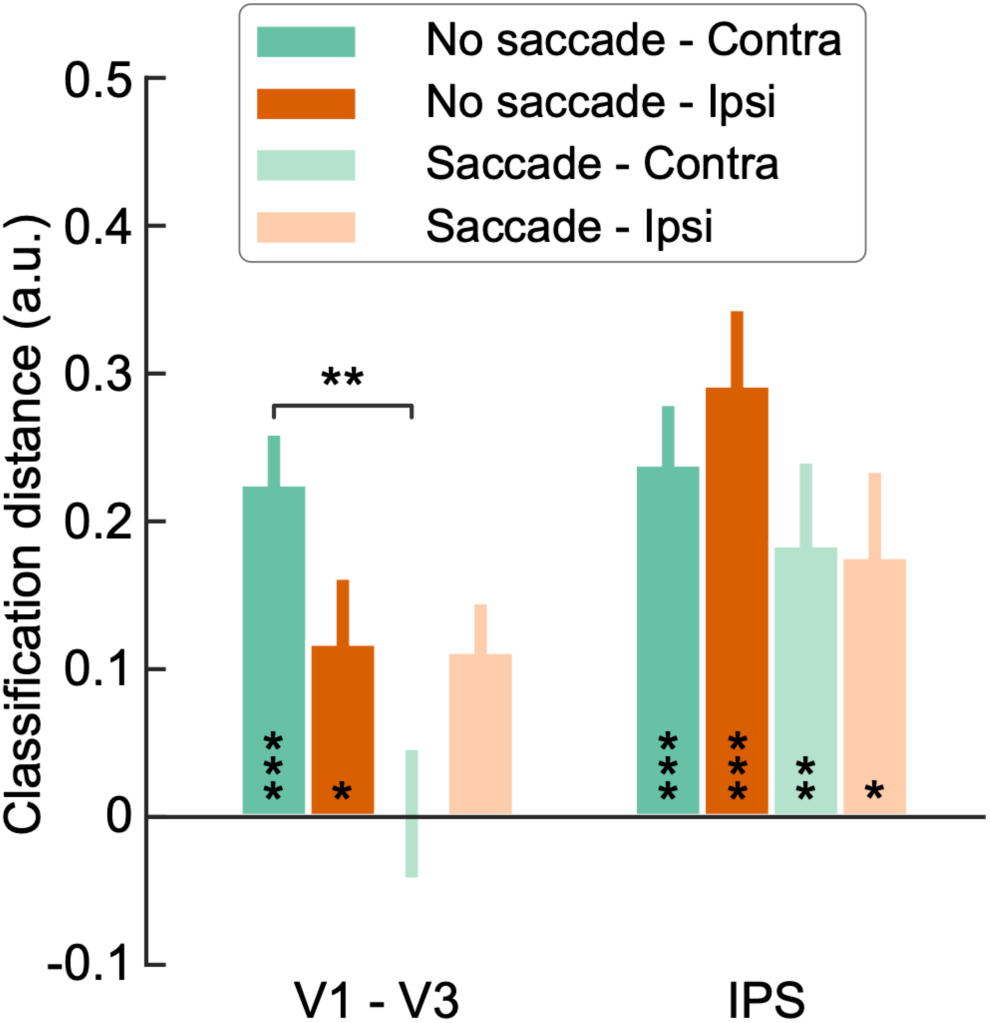
Orientation classification performance during the delay period in early visual cortex and intraparietal sulcus (IPS) using a leave-one-run-out cross-validation approach. In the combined early visual cortex (V1 – V3), both the contralateral and the ipsilateral early visual areas contained orientation information during the delay period in no-saccade condition. In the saccade condition, however, the orientation information significantly degraded in the contralateral hemispheres. Nevertheless, orientation information in the ipsilateral hemisphere after an eye movement was not higher than that before the saccade, suggesting no remapping of VWM in the early visual areas. In the IPS, the classifier could select the remembered orientation in both the contralateral and the ipsilateral hemisphere in the no-saccade trials, in line with the findings in early visual cortex. In the saccade condition, after the eye movement there was still a reliable orientation information in both the contralateral and ipsilateral IPS, suggesting a consistent VWM representation in the IPS. Error bars denote SEM. *No saccade – Contra*: No saccade condition, contralateral hemisphere; *No saccade – Ipsi*: No saccade condition, ipsilateral hemisphere; *Saccade – Contra*: Saccade condition, contralateral hemisphere; *Saccade – Ipsi*: Saccade condition, ipsilateral hemisphere. \**p* < 0.05; \*\**p* < 0.01; \*\*\**p* < 0.001.

Although orientation information in the contralateral hemisphere was degraded after the saccade, the remapping hypothesis of VWM predicts that orientation information should become stronger in the ipsilateral hemisphere after the eye movement. However, in contrast to this hypothesis, there was no difference in terms of orientation information in the ipsilateral hemisphere after the eye movement compared to when no saccade was made (t_(33)_ = 0.072, *p* = 0.9431, Cohen’s *d* = 0.017; BF_10_ = 0.184). Therefore, these results indicate that there is no remapping of VWM following eye movements in the early visual areas.

### Persistent VWM representations after an eye movement in the IPS

Our findings revealed that VWM representations in early visual cortex during maintenance were not remapped following eye movements. Bettencourt & Xu (2016) have recently shown that while remembered orientation information was degraded by irrelevant distractors in early visual cortex, VWM information remained available in the superior intraparietal sulcus (IPS). We sought to test whether we could observe similar results in our study. To this end, we also applied the leave-one-run-out cross-validation method to IPS. First, we found that both the contralateral (**Figure 2**, dark green bar; *t*_(33)_ = 4.451, *p* = 0.0001, Cohen’s *d* = 0.775; BF_10_ = 272.079) and ipsilateral IPS (**Figure 2**, dark orange bar; *t*_(33)_ = 5.075, *p* < 0.0001, Cohen’s *d* = 0.883; BF_10_ = 1444.41) contained information about the remembered orientation in no-saccade trials during delay period, as also observed in early visual cortex. However, in contrast to the early visual cortex, the remembered grating orientation could still be decoded from both the contralateral (**Figure 2**, light green bar; *t*_(33)_ = 3.13, *p* = 0.0036, Cohen’s *d* = 0.545; BF_10_ = 10.276) and ipsilateral (**Figure 2**, light orange bar; *t*_(33)_ = 2.6, *p* = 0.0138, Cohen’s *d* = 0.453; BF_10_ = 3.281) IPS after an eye movement, indicating a persistent VWM representations in IPS. These results also provided a neural evidence for supporting the preserved behavioral performance after a saccade.

Finally, in order to ensure that the results are not dependent on the a priori but arbitrarily chosen mask sizes of the ROIs, we repeated the analyses for ROIs of sizes ranging from 20 to 250 voxels in step of 10 voxels in the areas V1, V2 and V3, or ranging from 50 to 500 voxels in step of 20 voxels in the combined V1 – V3 and IPS (**Supplementary Figure 2**). Results were qualitatively identical to those mentioned above (**Figure 2** and **Supplementary Figure 1**) for almost entire range of ROI sizes, indicating that our results do not depend on an arbitrary ROI size but represent a robust pattern within the areas.

## Discussion

In the current study, we investigated whether VWM representations are remapped following eye movements and whether VWM information persists in the early visual cortex and parietal cortex after the execution of a saccade. We found robust encoding of maintained orientation information in the contralateral hemisphere of early visual cortex. However, this information significantly degraded upon making a saccade to the opposite hemifield and did not remap to the ipsilateral hemisphere in early visual cortex. This suggests that there is no robust representation of VWM representations across eye movements in the early visual cortex nor remapping of VWM representations. Additionally, although VWM representations in early visual cortex were impaired by the saccade during the retention period, this orientation information remained reliably present in the IPS even after execution of a saccade. These findings show that early visual cortex is susceptible to interference of eye movements during a working memory period, while IPS appears less sensitive to this interference.

### Spatially specific encoding of VWM information in early visual cortex?

The issue of how VWM information is stored in early visual cortex has been debated previously. While some researchers observed a retinotopically specific encoding of items into VWM (Pratte and Tong 2014), Ester and colleagues (2009) observed a non-spatially specific coding of VWM in early visual cortex. In their study, they could decode the orientation information in early visual areas both contralateral and ipsilateral to the stimulus location when participants were instructed to remember the orientation presented only in one visual hemifield. However, although the remembered grating was always located in one of the hemifields, participants could potentially use a strategy in which they memorized the orientation of the grating perifoveally instead of at its original location (Williams et al. 2008). Thus, it is unclear whether their result reflected the true effect of the spatially global representations or was caused by the lack of relevance of the spatial dimension in their task. To incentivize participants to form a spatially specific visual working memory, we inserted catch trials, in which the probe grating and the remembered grating were horizontally offset by 1.2 visual degree in each run. Participants were instructed to respond only when the probe location matched the location of the initially presented grating. Therefore, participants had to maintain the orientation of the gratings at the presented location in order to be able to successfully complete the task. Behavioral results confirmed that participants indeed remembered the orientation stimuli in a spatially specific way. Decoding performance in the early visual cortex (i.e., V1 – V3) showed a numerically larger amount of orientation information in contralateral than ipsilateral cortex, echoing earlier results (Pratte and Tong 2014) and suggesting a spatially specific code. However, this difference was not statistically reliable, and Bayesian analyses showed anecdotal evidence for a lack of difference (BF_10_ = 0.642). Thus, while our results may tentatively suggest some degree of retinotopic specificity of VWM encoding, our data preclude any strong conclusion.

### No remapping of VWM following eye movements

Attentional or feature based remapping of visual input just prior to a saccade has been well established from a variety of studies and is thought to maintain visual stability across eye movements. For instance, Rolfs et al. (2011) found that attention is predictively remapped to the future retinotopic location of an upcoming target, even before the execution of the eye movement. In the current experiment, we asked whether memory traces would also be updated with eye movements. However, we found no evidence that visual working memory representations were remapped following eye movements. At least two reasons should be emphasized to address the discrepancy in remapping between attention and VWM. First, to survive in a multifaceted and dynamically changing world, our attention needs to be selectively shifted to the most important stimuli that are goal-relevant. In order to be competent for such a complicated task, remapping of attention may be a crucial mechanism to prepare information in advance. In contrast, the main function of working memory is to temporarily retain information available for processing. While attention could be flexibly remapped to different locations to extract information, storing working memory information in one or several stationary regions could be potentially an efficient way for retrieval. Second, in line with the arguments of ‘activity-silent’ models of WM when compared to the persistent activity models of WM (Stokes 2015; Wolff et al. 2017), remapping of VWM after each eye movement could also be energetically expensive, especially if VWM is maintaining an up-to-date information of the external environment. Therefore, a stationary working memory representation could be an ideal way for ecologically maintaining and manipulating information in the brain.

### Parietal cortex may subserve VWM representations after saccades

In recent years, the parietal cortex has been repeatedly implicated in VWM maintenance (Todd and Marois 2004; Christophel et al. 2015; Ester et al. 2015; Bettencourt and Xu 2016), alongside the involvement of sensory areas (Harrison and Tong 2009; Serences et al. 2009; Emrich et al. 2013; Rademaker et al. 2019). Our results indicate that the early visual cortex is susceptible to the interference of eye movements during the delay period, potentially limiting the role of the early visual cortex in VWM. Similarly, Bettencourt and Xu (2016) observed that VWM representations in early visual cortex were destroyed by the presentation of distractors, while superior IPS exhibited persistent activity that was not impaired by the presentation of intervening distractor stimuli. In our experiment, the eye movements during the delay period potentially may also generate distracting new input to the visual system, which could disturb the VWM representations. Intriguingly, while VWM representations in early visual cortex were degraded following the eye movement, these representations were still maintained in IPS. Behaviorally, participants’ memory performance was indistinguishable between the saccade and no saccade trials. Therefore, it could be that the preserved behavioral performance we observed is supported by these IPS representations, rather than the early visual cortex representations. In a more recent study, Lorenc et al. (2018) found that the VWM information could be reliably maintained only in early visual cortex without distractors. However, these early visual cortex representations seem to transfer to the IPS representations when distractors were presented, which suggesting a flexible coding of VWM between the early visual cortex and the IPS depending on the task demands. This shows that VWM information maintained in visual cortex is susceptible to interference, in line with the idea that the early visual cortex may only play a limited role in VWM storage.

In conclusion, our results indicate that VWM representations in early visual cortex are degraded and not remapped after eye movements, but stably represented in parietal cortex. These results suggest a limited role of early visual cortex in VWM storage.

## Acknowledgements

This work was supported by The Netherlands Organisation for Scientific Research Vidi Grant 452-13-016 to F.P.d.L., the EC Horizon 2020 Program ERC Starting Grant 678286 “Contextvision” to F.P.d.L, the James S.McDonnell Foundation 220020373 to F.P.d.L, the China Scholarship Council (CSC; 201608330264) to T.H., and The Netherlands Organization for Scientific Research (Veni Grant No. 016.Veni.195.435) awarded to M.E.. We thank David Richter and Chuyao Yan for helpful comments and discussions of the manuscript.

## Author contributions

T.H. wrote the first draft of the paper; T.H., M.E., and F.P.d.L. edited the paper; T.H., M.E., A.V., and F.P.d.L. designed research; T.H. and A.V. performed research; T.H., M.E., and F.P.d.L. analyzed data.

## Supplementary information

**Supplementary Figure 1.**
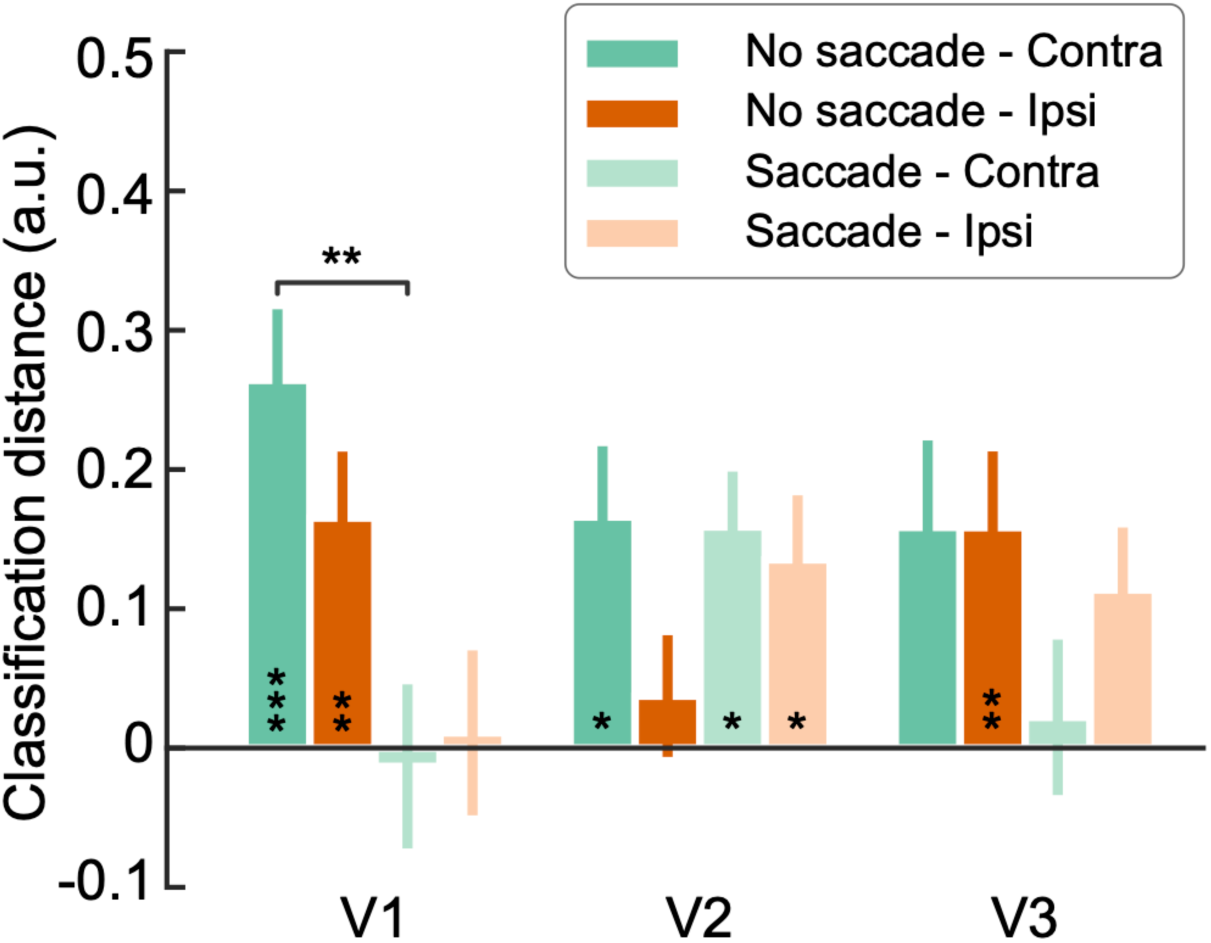
Orientation classification performance during the delay period in early visual areas separately. Same analysis as in Figure 2 but performed in V1, V2 and V3 individually. The same pattern was observed as in Figure 2. Specifically, in no-saccade trials, orientation information was present in the contralateral hemisphere of V1 (t_(33)_ = 3.701, *p* = 0.0008, Cohen’s *d* = 0.64; BF_10_ = 39.988), V2 (t_(33)_ = 2.460, *p* = 0.0193, Cohen’s *d* = 0.43; BF_10_ = 2.48), and V3 (t_(33)_ = 1.892, *p* = 0.0673, Cohen’s *d* = 0.33; BF_10_ = 0.901), and also in the ipsilateral hemisphere of V1(t_(33)_ = 2.883, *p* = 0.0069, Cohen’s *d* = 0.502; BF_10_ = 5.945) and V3 (t_(33)_ = 3.147, *p* = 0.0035, Cohen’s d = 0.548; BF_10_ = 10.688), but not V2 (t_(33)_ = 0.612, *p* = 0.5447, Cohen’s *d* = 0.107; BF_10_ = 0.219). In saccade trials, however, the classifier was not able to distinguish between grating orientations in the contralateral hemisphere of the V1 (t_(33)_ = -0.174, *p* = 0.8627, Cohen’s *d* = 0.03; BF_10_ = 0.186) and V3 (t_(33)_ = 0.313, *p* = 0.7562, Cohen’s *d* = 0.054; BF_10_ = 0.192), but not V2 (t_(33)_ = 2.542, *p* = 0.0159, Cohen’s *d* = 0.443; BF_10_ = 2.917), while the orientation information remained stable in the ipsilateral hemisphere of the V2 (t_(33)_ = 2.306, *p* = 0.0276, Cohen’s *d* = 0.395; BF_10_ = 1.848) and V3 (t_(33)_ = 2.028, *p* = 0.0507, Cohen’s *d* = 0.348; BF_10_ = 1.128), but not V1 (t_(33)_ = 0.132, *p* = 0.8959, Cohen’s *d* = 0.023; BF_10_ = 0.185). The orientation information in the contralateral hemisphere of the V1 was significantly degraded by an eye movement (t_(33)_ = 2.765, *p* = 0.0093, Cohen’s *d* = 0.647; BF_10_ = 4.612). Error bars denote SEM. *No saccade – Contra*: No saccade condition, contralateral hemisphere; *No saccade – Ipsi*: No saccade condition, ipsilateral hemisphere; *Saccade – Contra*: Saccade condition, contralateral hemisphere; *Saccade – Ipsi*: Saccade condition, ipsilateral hemisphere. \**p* < 0.05; \*\**p* < 0.01; \*\*\**p* < 0.001.

**Supplementary Figure 2.**
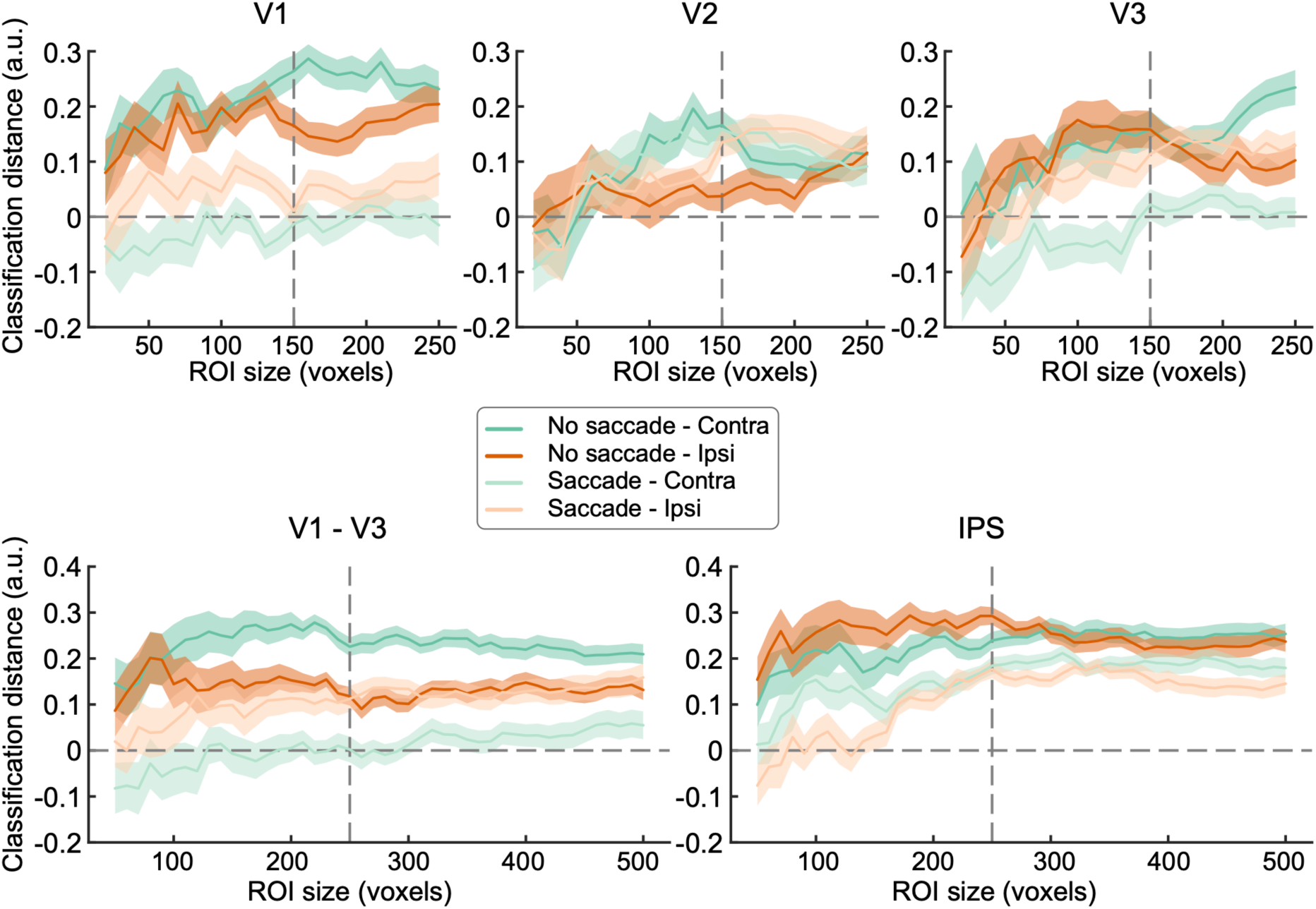
Orientation classification performance during the delay period over a range of ROI sizes using a leave-one-run-out cross-validation method. Same analysis as in Figure 2 but performed over a wide range of ROI sizes. In areas V1, V2, and V3, we rerun the analysis for ROIs of sizes ranging from 20 to 250 voxels in step of 10 voxels, while in areas V1 – V3 and IPS, the analysis was repeated for ROIs of sizes ranging from 50 to 500 voxels in step of 20 voxels. The dashed vertical line is at the predefined ROI of 150 voxels in areas V1, V2, and V3, or 250 voxels in areas combined V1 – V3 and IPS. The same pattern of effects was found for almost entire range of ROIs. Interestingly, in the combined V1 – V3, while WM information in the contralateral hemisphere was reduced after an eye movement (dark green line vs. light green line), it was persistently presented in the ipsilateral hemisphere for trials with and without a saccade (dark orange line vs. light orange line) over all range of ROI sizes. In IPS, however, no matter in the saccade or no-saccade condition, both the contralateral and ipsilateral hemispheres showed consistent orientation information over the range of ROIs, suggesting a robust VWM representation in this region.

### Supplementary table

**Supplementary Table 1.**
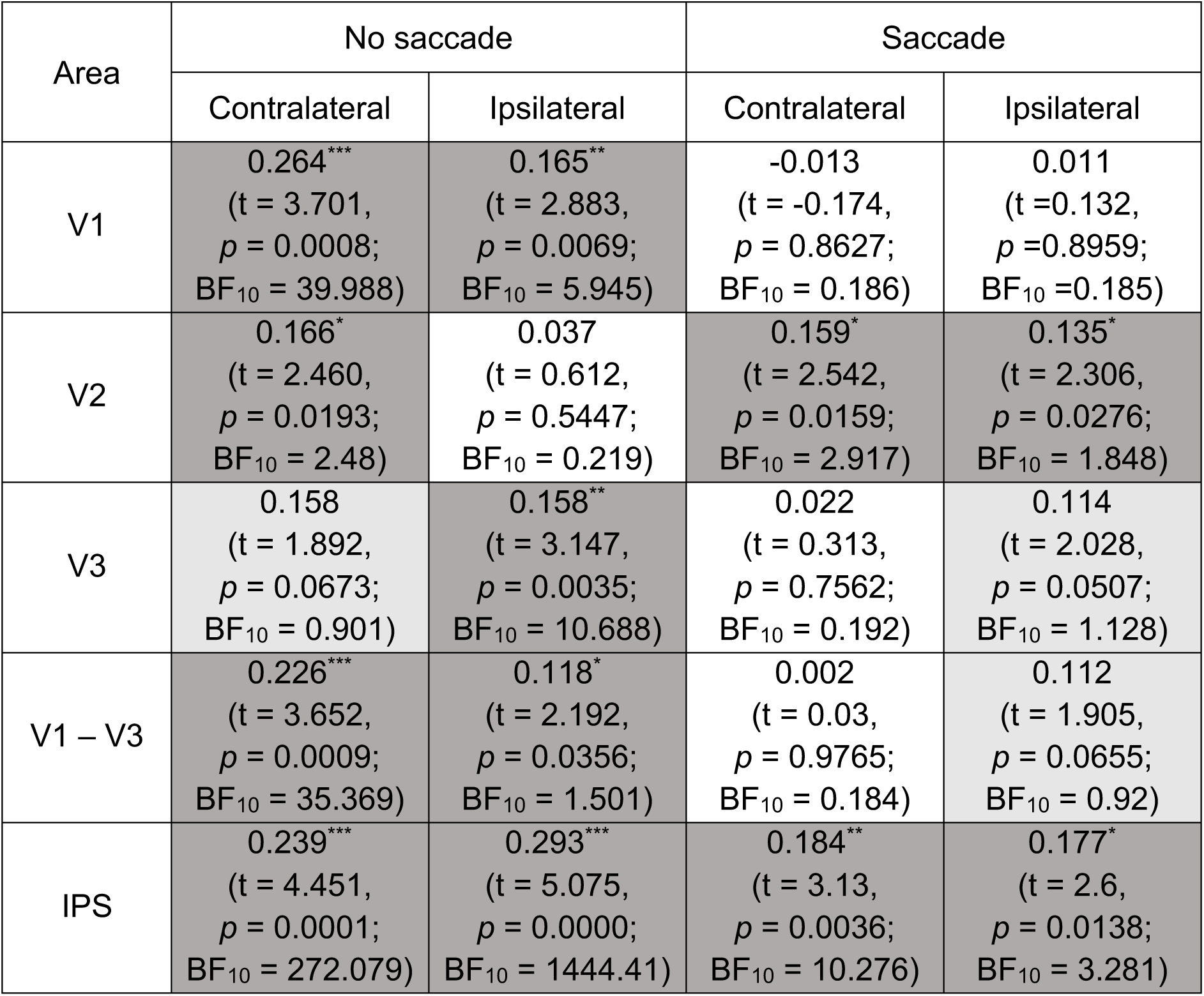
Decoding distance over 7.2 – 10.8 seconds after the onset of the delay period for each ROI and post-hoc t-test of significant differences (see Figure 2). Dark grey cells indicate that there was a significant effect of saccade and hemisphere condition in that ROI, light grey cells represent the effect only reached a marginal significance, while cells indicate that there was no significant effect in that ROI. \**p* < 0.05; \*\**p* < 0.01; \*\*\**p* < 0.001.

| Area | No saccade |  | Saccade |  |
| --- | --- | --- | --- | --- |
|  | Contralateral | Ipsilateral | Contralateral | Ipsilateral |
| V1 | 0.264***<br>($t = 3.701$,<br>$p = 0.0008$;<br>$BF_{10} = 39.988$) | 0.165**<br>($t = 2.883$,<br>$p = 0.0069$;<br>$BF_{10} = 5.945$) | -0.013<br>($t = -0.174$,<br>$p = 0.8627$;<br>$BF_{10} = 0.186$) | 0.011<br>($t = 0.132$,<br>$p = 0.8959$;<br>$BF_{10} = 0.185$) |
| V2 | 0.166*<br>($t = 2.460$,<br>$p = 0.0193$;<br>$BF_{10} = 2.48$) | 0.037<br>($t = 0.612$,<br>$p = 0.5447$;<br>$BF_{10} = 0.219$) | 0.159*<br>($t = 2.542$,<br>$p = 0.0159$;<br>$BF_{10} = 2.917$) | 0.135*<br>($t = 2.306$,<br>$p = 0.0276$;<br>$BF_{10} = 1.848$) |
| V3 | 0.158<br>($t = 1.892$,<br>$p = 0.0673$;<br>$BF_{10} = 0.901$) | 0.158**<br>($t = 3.147$,<br>$p = 0.0035$;<br>$BF_{10} = 10.688$) | 0.022<br>($t = 0.313$,<br>$p = 0.7562$;<br>$BF_{10} = 0.192$) | 0.114<br>($t = 2.028$,<br>$p = 0.0507$;<br>$BF_{10} = 1.128$) |
| V1 – V3 | 0.226***<br>($t = 3.652$,<br>$p = 0.0009$;<br>$BF_{10} = 35.369$) | 0.118*<br>($t = 2.192$,<br>$p = 0.0356$;<br>$BF_{10} = 1.501$) | 0.002<br>($t = 0.03$,<br>$p = 0.9765$;<br>$BF_{10} = 0.184$) | 0.112<br>($t = 1.905$,<br>$p = 0.0655$;<br>$BF_{10} = 0.92$) |
| IPS | 0.239***<br>($t = 4.451$,<br>$p = 0.0001$;<br>$BF_{10} = 272.079$) | 0.293***<br>($t = 5.075$,<br>$p = 0.0000$;<br>$BF_{10} = 1444.41$) | 0.184**<br>($t = 3.13$,<br>$p = 0.0036$;<br>$BF_{10} = 10.276$) | 0.177*<br>($t = 2.6$,<br>$p = 0.0138$;<br>$BF_{10} = 3.281$) |

